# Spatial MIST Technology for Rapid, Highly Multiplexed Detection of Protein Distribution on Brain Tissue

**DOI:** 10.1101/2022.01.11.475923

**Authors:** Revanth Reddy, Liwei Yang, Jesse Liu, Zhuojie Liu, Jun Wang

## Abstract

Highly multiplexed analysis of biospecimens significantly advances the understanding of biological basics of diseases, but these techniques are limited by the number of multiplexity and the speed of processing. Here, we present a rapid multiplex method for quantitative detection of protein markers on brain sections with the cellular resolution. This spatial multiplex *in situ* tagging (MIST) technology is built upon a MIST microarray that contains millions of small microbeads carrying barcoded oligonucleotides. Using antibodies tagged with UV cleavable oligonucleotides, the distribution of protein markers on a tissue slice could be “printed” on the MIST microarray with high fidelity. The performance of this technology in detection sensitivity, resolution and signal-to-noise level has been fully characterized by detecting brain cell markers. We showcase the codetection of 31 proteins simultaneously within 2 h which is about 10 times faster than the other immunofluorescence-based approaches of similar multiplexity. A full set of computational toolkits was developed to segment the small regions and identify the regional differences across the entire mouse brain. This technique enables us to rapidly and conveniently detect dozens of biomarkers on a tissue specimen, and it can find broad applications in clinical pathology and disease mechanistic studies.

## INTRODUCTION

Human tissue is associated with a complex architecture that houses a variety of spatially distributed cell types. Each cell is a relatively separate unit within a neighborhood executing its functions, and its intricate machinery involves numerous proteins, RNAs and small molecules in networks. Thus, fully understanding of the cellular status and phenotypes normally requires highly multiplexed or even omics technologies to profile the cellular contents.^1, 2^ Proteins predominate for representing cell phenotypes, drug targets, clinical biomarkers, signaling networks, transcriptional factors, functional readouts of proliferation, cell cycle status, metabolism regulation and apoptosis makers.^3–5^ Thus, it is necessary to have highly multiplexed technologies to analyze protein contents at the cellular level. Conventional immunohistochemistry (IHC)-based techniques only analyze 1-4 protein markers of interest at a time, which could not capture the complexity of the cells in a tissue architecture. Mass spectrometry-based proteomic techniques often lose the spatial-anatomical context and regional information or lack single-cell sensitivity.^6–9^

Many efforts have been made to enhance the multiplexity of IHC technique since it is simple, inexpensive, and assessable to common users. Fluorescence microscopy enables quantitative analysis of multiple proteins through spectrally resolved fluorophores, and up to seven fluorophores have been reported for multispectral imaging.^10^ Raman-active dyes could overcome that limit by analyzing up to 20 antigens due to broad wavelength range and narrow vibrational peaks of the dyes,^11, 12^ which, however, needs a special Raman microscope not available to many users. Multiplex ion beam imaging can image 30+ proteins at once with the subcellular resolution but it still needs a special instrument.^13^ An alternative approach is cyclic immunofluorescence (IF) staining to enhance multiplexity by reiteratively labeling the same tissue with fluorescent dyes and registering the images, so the same cells are stained for multiple times.^14–20^ This method can conveniently analyze dozens of proteins with even 20 rounds of labeling, antibody dissociation (or photobleaching) and re-labeling.^21^ However, cyclic IF and the variants may unpractically take >2 days to analyze 30+ markers, and the repeated processing of the same tissue could cause antigen loss and structural damage.^22, 23^ Besides, intrinsic autofluorescence of brain tissue could interfere with the IF-based detection.

In this report we describe a rapid, one-shot approach that could analyze dozens of protein markers on a tissue slice within 2 h from fixation to acquirement of multiplexed signal, using a common fluorescence microscope. This method is based on IHC where the staining on a tissue slice is spatially transferred to a multiplex *in situ* tagging (MIST) microarray. We have fully characterized this spatial MIST technology by examining the sensitivity, spatial resolution and signal-to-noise level. The spatial MIST was validated by detecting 1 or 5 markers simultaneously on a mouse brain slice. Definition of pseudocells permits detailed analysis of regional differences of brain slices. Further application by codetection of 31 markers led to the discovery of brain sub-regions represented by individual proteins and their spatial proximity distances. The spatial MIST is simple and easy to reproduce with high robustness, and it only requires a common fluorescence microscope for imaging. It is expected to advance information-rich pathological analysis of clinical samples in the future as well as mechanistic studies of brain diseases.

## EXPERIMENTAL SECTION

### MIST array preparation and characterization

Polystyrene microbeads (Thermo fisher) were coated with single-stranded DNA (ssDNA) oligonucleotides first before patterning of a monolayer. Amine-polystyrene microbeads (2 μm; Life Technologies) were coated with poly-L-lysine (PLL; Ted Pella) to amplify the amount of amine groups on the surface before DNA oligonucleotide anchoring. Amine-microbeads were treated with 10 mM bis(sulfosuccinimidyl)suberate (BS3; Pierce) crosslinker in phosphate buffer saline (PBS; pH 7.4) for 20 min. After washing by Milli-Q water two times, PLL was mixed with the activated microbeads in a pH 8.5 PBS solution for 4 h. The PLL-coated microbeads were rinsed by PBS with 0.05% Tween 20 three times and were further reacted with amine-ended ssDNA at 300 μM and 2 mM BS3 for 4 h. The microbeads were then thoroughly washed and resuspended to the original concentration for further use.

Microbeads of one or multiple types of equal portion with oligo DNA coating were washed first with DI water and sonicated to separate the aggregates. The microbeads were concentrated to be >50% w/v by centrifugation and then deposited onto a cleanroom adhesive tape (VWR) attached on a glass slide. The array was sonicated for 1 min to remove the excess layers of microbeads on a monolayer, which was validated by examination under a microscope. The MIST array can be stored dry at 4°C for weeks.

To evaluate the quality of the MIST array, a cocktail of Cy5-tagged complementary oligo DNAs at 200 nM in tris buffer with 0.05% tween 20 (TBST) was applied to the MIST array and incubated for 1 h. The array was washed for 5 times by TBST and imaged under a fluorescence microscope (Nikon Ti-2). The fluorescence intensities of each microbead of every type were quantified by a MATLAB program developed in the lab. The sensitivity of the MIST array was measured by the use of various concentrations (1 pM, 10 pM, 100 pM, 1 nM and 10 nM) of complementary oligo DNAs tagged with Cy5 (Integrated DNA Technologies). The fluorescence intensities of microbeads were quantified to fit in a logistic function to quantify the detected oligo DNAs in the solution. Limit of detection (LOD) was calculated by the background signal plus three times of its standard deviation on the fitting curve.

### Preparation and purification of UV-cleavable, biotinylated DNA-antibody conjugates

Conjugation of biotinylated oligo DNAs and antibodies was achieved through DBCO-azide click chemistry reaction.^24^ The antibodies were concentrated to 1 mg/mL first using 10K MWCO centrifuge filters (Amicon; Thermo fisher) and were purified by 7K MWCO Zeba spin desalt columns (Thermo fisher) while the pH of the solution was adjusted to 8.0. 50 μL antibodies were then modified with UV-cleavable azido-NHS ester (Click Chemistry Tools) at 1:15 molar ratio for 2 h. Meanwhile the solutions of the amine-ended, biotinylated DNAs at 200 μM were also adjusted to pH 8.0 and conjugated with UV cleavable DBCO-NHS ester (Click Chemistry Tools) at 1:20 molar ratio for 2 h. The modified antibodies and oligo DNAs were purified and suspended in pH 7.4 PBS using 7K MWCO Zeba spin desalting column to remove excess chemicals, and they were mixed for crosslinking overnight at room temperature. The conjugates were further purified on a FPLC workstation (AKTA; Bio-Rad) with Superdex 200 gel filtration column at 0.5 ml/min. The collected products were concentrated to 0.3-0.5 mg/mL and stored at 4°C for further use. To determine the conjugation efficiency, the absorbance spectrum of a UV-cleavable conjugate was measured by a Nanodrop spectrophotometer (Thermo fisher), and the degree of labeling was calculated as previously reported.^25^

### Brain tissue slice and immunofluorescence staining

The mouse C57 hippocamp/thalamus/hypothal coronal frozen section was made of freshly harvested mouse brain (Zyagen). The tissue slices were sectioned at a thickness of 7-10 μm and mounted on positively charged slides. The brain slices were fixed by 4% formaldehyde (Thermo fisher) for 10 min, followed with permeabilization by 0.1% Triton X100 for 10 min. The slices were blocked by 5% goat serum (Cell Signaling) in PBS for 1 h, washed by the same blocking buffer, and stained with the primary antibodies at the recommended concentrations by the vendors for 1 h. The secondary antibodies tagged with Alexa Fluor 647 (Invitrogen) at 5 μg/mL were added to the slices and incubated for 1 h before three-times washing and imaging.

### Spatial MIST detection and characterization

The brain slice was fixed and permeabilized as described above before incubating with a blocking buffer containing 5% goat serum, 100 μg/mL sheared Salmon sperm DNA (Thermo fisher) and 0.05% tween 20 in PBS for 45 min. Then a single conjugate, or a mixture of conjugates, that was diluted 100 times in the blocking buffer was applied to the brain slice and incubated for 1 h in dark. Meanwhile a MIST array carrying the complementary DNA coated microbeads was blocked by a TBST buffer for 10 min. After washing of the slice and the MIST array for three times, they were mated and clamped tightly by magnetic force. The whole setup was exposed to a UV light (365 nm; Thorlabs CS2010 UV Curing LED System) for 5 min. After separation in a PBS buffer, the brain slice was stained by secondary antibodies with the procedure described above while the MIST array was thoroughly washed by PBS buffer. Fluorescence images of the tissue were taken using a fluorescence microscope. For detection of signal on the MIST array, the array was incubated with 10 μg/mL streptavidin-Alexa Fluor 647 (Invitrogen) in PBS solution containing 3% BSA (Sigma) for 15 min. The array slide was washed by the same buffer for 5 times before microscopy imaging and scanning.

To analyze the influence of UV exposure time on the detection, the tissue slices within the clamp were exposed with the UV light for 2 min, 5 min, 10 min and 20 min when only FOX3 was detected. The MIST arrays were washed with PBS buffer immediately after separation from the tissue slices. The fluorescence intensities of each single cell in the prefrontal cortex area were quantified. The average intensity of cells was divided by the average intensity of the same area to generate the signal to noise ratio.

The detection sensitivity was measured by the capability of detecting complementary oligos. Cy5-labeled complementary oligo solutions at various concentrations (1 pM, 10 pM, 100 pM, 1 nM and 10 nM) were applied to the array and incubated for 1 h, followed by PBS washing and imaging under a microscope. The intensities of each microbead were quantified by ImageJ. The limit of detection (LOD) for the oligos in solution were calculated by the background signal plus three standard deviations after extrapolating the logistic fitting curve.

### MIST array decoding process

After detection, the entire MIST array was scanned with a high NA 20X objective to record the protein signals on each microbead. The MIST array was processed by 150 μL 0.5 M NaOH solution for 1 min to dissociate double stranded DNAs on the microbeads, before thoroughly washing with saline-sodium citrate buffer (SSC) for five times. A cocktail of complementary DNA tagged with various fluorophores at 200 nM in a hybridization buffer (40% formamide and 10% dextran sulfate in SSC buffer) for each was applied to the MIST array for 1 h at room temperature. Then the array was washed for three times by SSC buffer and subsequently imaged under a fluorescence microscope, to complete “Cycle 1” of the decoding procedure. The array was scanned by the same approach as in the protein detection. The second and third cycles, denoted as “Cycle 2” and “Cycle 3”, were executed by the same procedure except that the cocktails of complementary DNA-fluorophores are different. For the detection of 5 proteins, only two cycles were executed in decoding. All the images from protein detection to decoding cycles were registered together to determine the order of fluorescent color change for each microbead, while the order was predesigned for each type of oligo DNA microbead or protein type.

### MIST array data requisition and image registration

A Nikon Ti2 inverted fluorescence microscope equipped with a motorized stage was used to automatically take images for the MIST arrays and the brain slices. A bright field image in high contrast was taken in each area to recognize the location of individual microbeads. For the decoding image suite of each cycle, the fluorescent colors for Alexa Fluor 488, Cy3, Cy5 and Alexa Fluor 750 were automatically imaged by a wavelength switcher. All the images were saved as the format of 16-bit ND2 in the Nikon software (NIS-Elements AR Microscope Imaging Software) and then were passed to MATLAB program for further registration and processing.

A MATLAB program was developed for data requisition and image registration. For each cycle, decoding images were registered with the bright field images. Once the decoding images were aligned, contrast was enhanced by equalizing intensity histograms for each decoding image. The locations and median intensity of individual microbeads were detected. Each microbead was assigned a color for each decoding cycle depending on the highest intensity fluorescent color in the associated decoding images. To reduce the impact of background noise on color detection, the microbeads were only assigned colors if the intensity of each color was over 2 times the intensity of the background. Each microbead was then labeled with a specific protein depending on the sequence of colors for each cycle. Image registration error was approximated by measuring the mean translational distance of the corresponding points that were improperly transformed during image registration. Corresponding points were defined as pixels containing the same intensity values in both the fixed and moving images. In ideal image registration, these points should not be displaced after registration. The mean displacement of all corresponding points after registration was calculated for each decoding cycle in all images.

### Image reconstruction

Once the position, intensity, and identity (ID) for each microbead were determined, the full image was reconstructed by stitching together individual image areas. The spatial distribution of a selected protein type was visualized by retaining only the microbead signal of this protein and setting all other regions as dark background on the stitched image. 2.5% of all edges of the individual images was also set as dark background since image registration might have left out edge microbeads. To increase visibility of individual protein distributions, an averaging filter was applied to all images.

### Pseudocells

Pseudocells were generated in each image to assess segments of various local regions. First, each image was cropped by 2.5% on each side to reduce the impact of unidentified beads located on the edges of each image. The intensity and distance of the nearest microbeads of the same type were measured to determine the size of pseudocells. A histogram of distances binned at 1 μm was then generated to determine the likelihood of the pseudocell containing each type of protein. In principle, each pseudocell should contain all type of microbeads with >95% probability. The size of a pseudocell was determined as a 20 μm by 20 μm unit in a grid across the entire image for the 31-plex assay. For the multiplexed detection of 5 proteins, the dimensions of each pseudocell were 6 μm by 6 μm instead.

### Clustering and statistics

Uniform manifold approximation and projection (UMAP) was used for dimension reduction and visualization of pseudocell clusters using the R package “Seurat” with the UMAP parameters (dims=1:26, min.dist=0.1, spread=2, n.neighbors=50).^26^ In the preprocessing step, principal component analysis (PCA) removed the significant outliers including protein 19 and 20. The background and the regions without signal were further removed through filtering, where the threshold for each protein was set at the mean intensity of lowest 1 percentile of all pseudocells plus three standard deviations. The pseudocells with lower than 20% of proteins detectable were also dropped. Then they were log2 normalized and clustered by using the Python packages NumPy and pandas to visualize the UMAP clusters.^27^

The spatial proximity distance between pairs of subtypes in heatmap was created by first assigning the pseudocells with a cluster number determined by the Seurat clusters from the UMAP plot. Global pseudocell coordinates was used to quantify the spatial distance between a pseudocell cluster and another pseudocell cluster by finding the minimum nearest neighbor distances between the first cluster’s pseudocell and all of the second cluster’s pseudocells for each of the first cluster’s pseudocell. The average of all the minimum nearest neighbor distance values was then taken as the final value of spatial distance. This methodology was repeated for each pair of pseudocell cluster, including switching the cluster pair order between first cluster and second cluster. From the resulting matrix, the matrix values (average minimum nearest neighbor distances) were scaled, normalized, and multiplied by the maximum of the original matrix’s value and then used to construct the heatmap using the R package lattice and its color scale was constructed using the R package color space.^28, 29^

## RESULTS AND DISCUSSION

The spatial MIST technology permits ‘printing’ of the local molecules of a tissue slice to the MIST array (Figure 1a). The essential components of the technology are a super compact barcode oligonucleotide DNA array and custom-designed, UV-cleavable complementary oligonucleotide-antibody conjugates. The former contains ~25,000,000 barcode DNA oligonucleotide carrying microbeads in a 1 cm x 1 cm area on a glass slide. The latter allows for conversion of protein detection to DNA detection and use DNA techniques for multiplexed analysis.^30, 31^A mouse brain coronal section was stained with a mixture of oligonucleotide-antibody conjugates which contains photocleavable linkers. When the stained tissue was mated with a MIST array, the oligos were released upon UV exposure and were captured locally by the complementary oligo DNAs on the MIST array. Figure 1b shows the detection of FOX3 protein on a tissue slice by immunofluorescence (IF) staining and on the MIST array of the same slice. The ‘printed’ image matches the one on the tissue in high fidelity where the single cells can be distinguished. Interestingly, the autofluorescence on the tissue sample, which is a common issue in IF, is completely gone on the MIST array, as bright patches of signals were not observed on the MIST array image.

**Figure 1.**
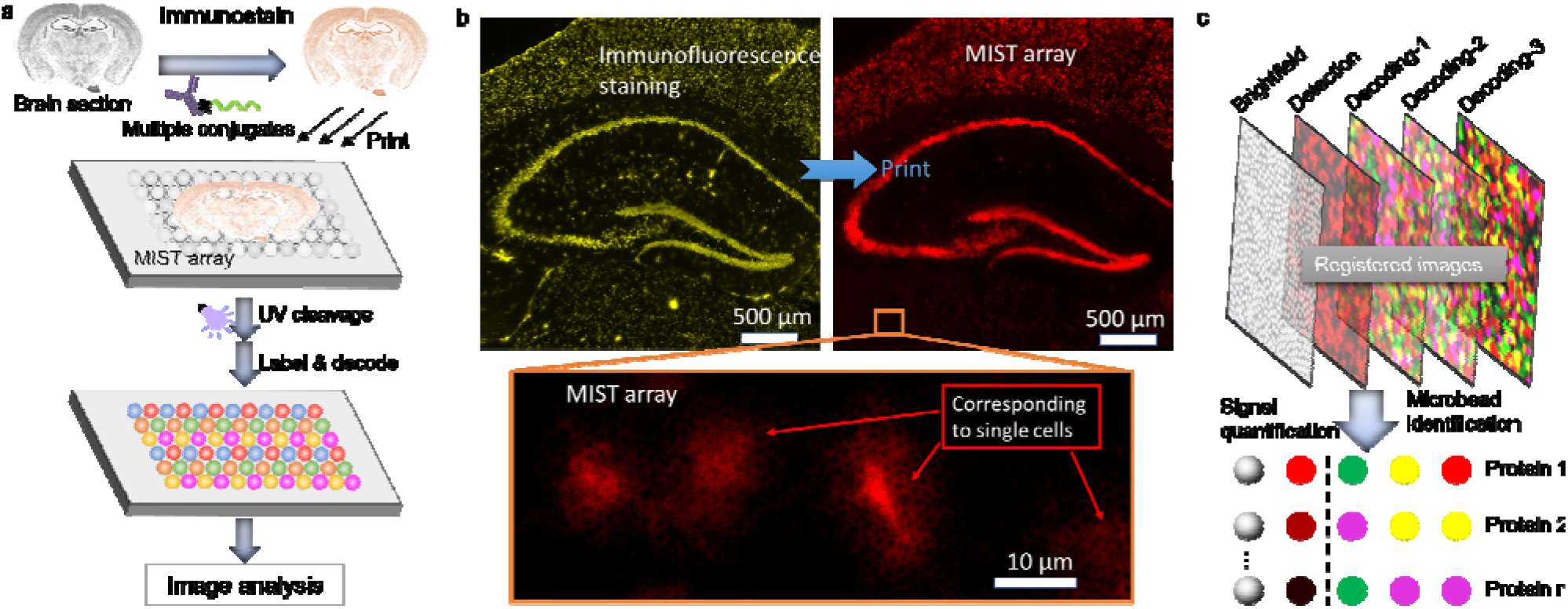
Principle of spatial MIST for multiplexed detection of proteins. (a) Overview of the assay process. (b) Distribution of FOX3 protein by immunofluorescence staining and MIST array on the same brain slice. The zoom-in image below shows the individual neurons in the thalamus region of the brain sample. (c) Assignment of protein identities to each microbead through reiterative labeling and dissociation of complementary DNA oligos tagged with fluorophores while the protein detection signal is quantified on the individual microbeads. The location of microbeads is recorded for reconstruction and data analysis.

For multiplexed detection, we employed a decoding process through reiterative staining and dissociation to assign the identities to each microbead and quantify the expression of each protein. The brightfield image was used to locate individual microbeads. It was registered with protein detection image and three decoding images so all of them were aligned with 0.24 μm error in average, which is much less than 2 μm of the microbead diameter. Thus, each microbead would have a full set of data including protein detection signal and four fluorescent colors where each sequence of colors is uniquely corresponding to one type of oligonucleotides or proteins. The multiplexity of this technique exponentially increases the cycle number or the color number. In our study, 4 colors with 2 cycles can detect up to 16 types of proteins, and one more cycle leads to detection of up to 64 proteins. In our previous report without spatial information, the MIST array has been applied to detect up to 50 proteins per cell each time.^32^

The performance of spatial MIST has been characterized by the analysis of detection sensitivity, signal to noise ratio (SNR) and spatial resolution. We quantified the sensitivity of the MIST array by measuring the fluorescence signals of various concentration of complementary oligos in the solution that simulates the release of oligos from the tissue (Figure 2a). The panel of 31 oligos shows high sensitivity with mean detection limit at 4.0 pM, which is partially attributed to the high loading of oligos on the microbead surface through PLL mediation. Since two UV cleavable linkers were designed between an oligo and the antibody, >99% of oligos has been released.^32^ Thus, the detection on the MIST array quantitatively reflects on the amount of antibodies binding to their targets on the tissue. Notably, most IF or immunohistochemistry-based methods don’t quantify the amount of target proteins since not all protein epitopes are achievable for antibody recognition and binding. The SNR of the spatial MIST assay was assessed with various UV exposure durations of 2 min, 5 min, 10 min, and 20 min (Figure 2b). The fluorescence intensities of microbeads of single cells on the MIST array in the cortical region were measured, and the mean value was divided by the mean intensity of the non-cell area of the same region (see zoom-in image on Figure 1b). It was found that 5 min exposure time resulted in the best signals of single cells and also the best SNR. Longer time is not beneficial as both signals of single cells and SNR gradually decrease after 5 min. That is because the released oligos diffused farther with longer UV exposure time and incubation time.

**Figure 2.**
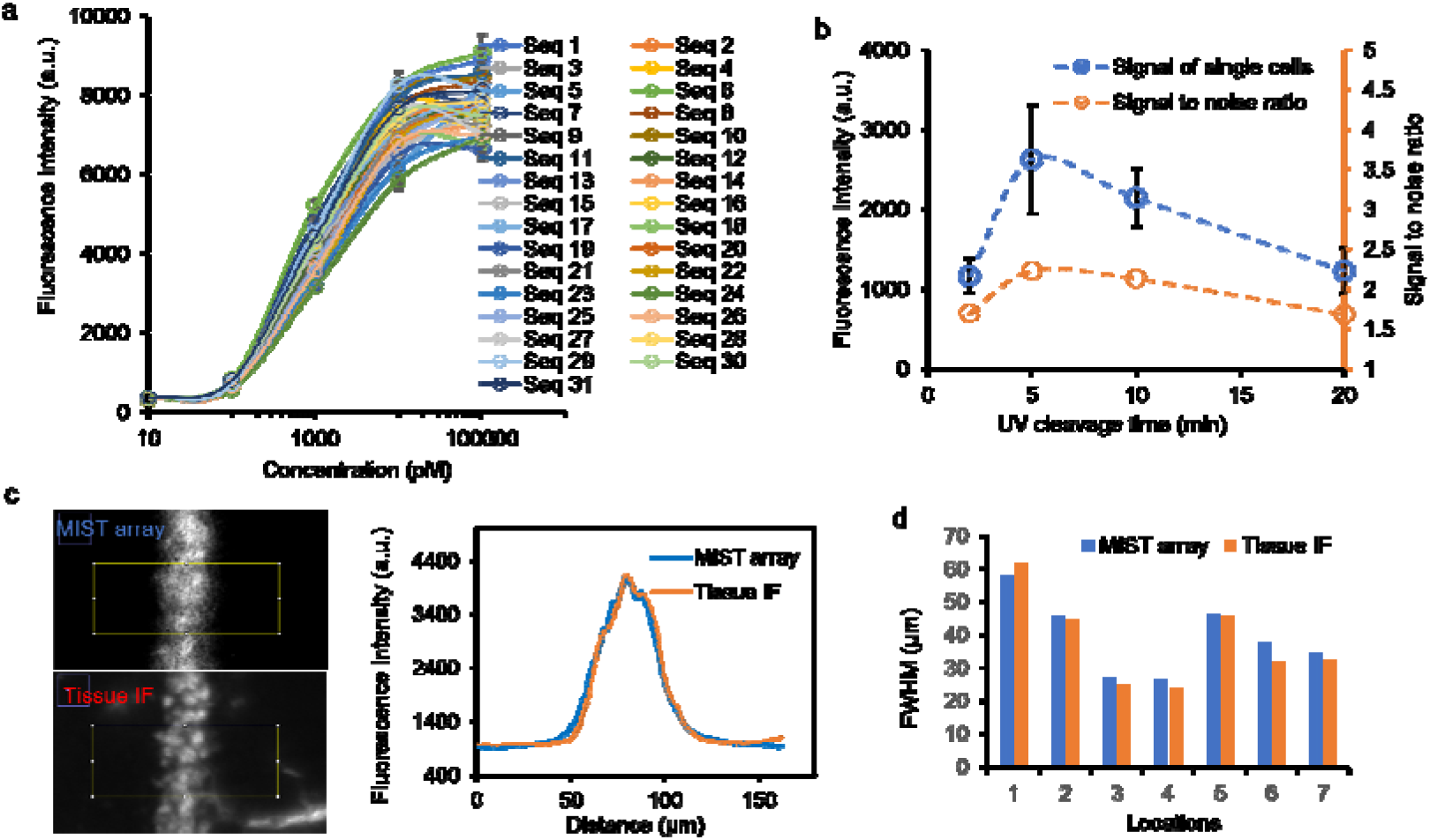
Characterization of detection by the spatial MIST technology. (a) Sensitivity of the MIST array for multiplexed detection of 31 DNAs. Calibration curves for each DNA were generated using complementary DNA oligos tagged with Alexa Fluor 647 for hybridization to the DNA oligos on the MIST array at various concentrations. (b) Quantified fluorescence signals of individual cells in the isocortex region of the coronal section. The sample was incubated for 2 min, 5 min, 10 min and 20 min with UV light on during spatial MIST assay. The error bars for the signal curve represent variation of 150 cells. The curve of signal-to-noise ratio is superimposed on the right y axis. (c) Comparison of the selected area from tissue IF and MIST array images when FOX3 is detected. The ROI is shown in rectangle in the CA1 region of the brain coronal section while the fluorescence intensity distribut on curves for the two ROIs are shown on the right chart. (d) Quantified FWHM of 7 selected ROIs on the CA1 region from MIST array and tissue IF images.

We examined the single neurons of the cortical region to characterize the spatial resolution of detection. The full width at half maximum (FWHM) of single cells on the MIST array is 86.9±12.6% of the same cells on the tissue, indicating sharper, more distinctive cell signals detected by the MIST. It was found that the profiles of individual cells on the MIST are slightly more spreading out on the edges than those of the same immunostained cells, but the centers are brighter. Thus, once the cells were adjacent to each other (Figure 2c), the individual cells were less distinctive on the MIST array. However, the overall distribution of the CA1 stripe is similar in two cases. We also analyzed the various regions of the CA1 stripe and found those profiles of the MIST array and the tissue IF are always similar with <5% deviation (Figure 2d).

The spatial MIST was applied for multiplexed detection of 5 brain cell markers including FOX3, GFAP, TBR1, CUT1 and CD11b. The codetection of all 5 markers and the decoded images for individual proteins are shown in Figure 3a. In the decoded images, only the microbeads for detecting a particular protein remained and all the others were masked in dark. Figure 3b shows the hippocampal region of the mouse brain where FOX3 detection reveals the distribution of neuronal cells. The heatmap is corresponding to the shaded area of the CA1 region, and it clearly shows the stripe of neurons as expected. To assist further analysis, the entire stitched brain image was segmented into pseudocells of 6 μm by 6 μm for each. This size was chosen because at least one copy of microbeads of any of 5 types should be present in the pseudocell of all 5 proteins.

**Figure 3.**
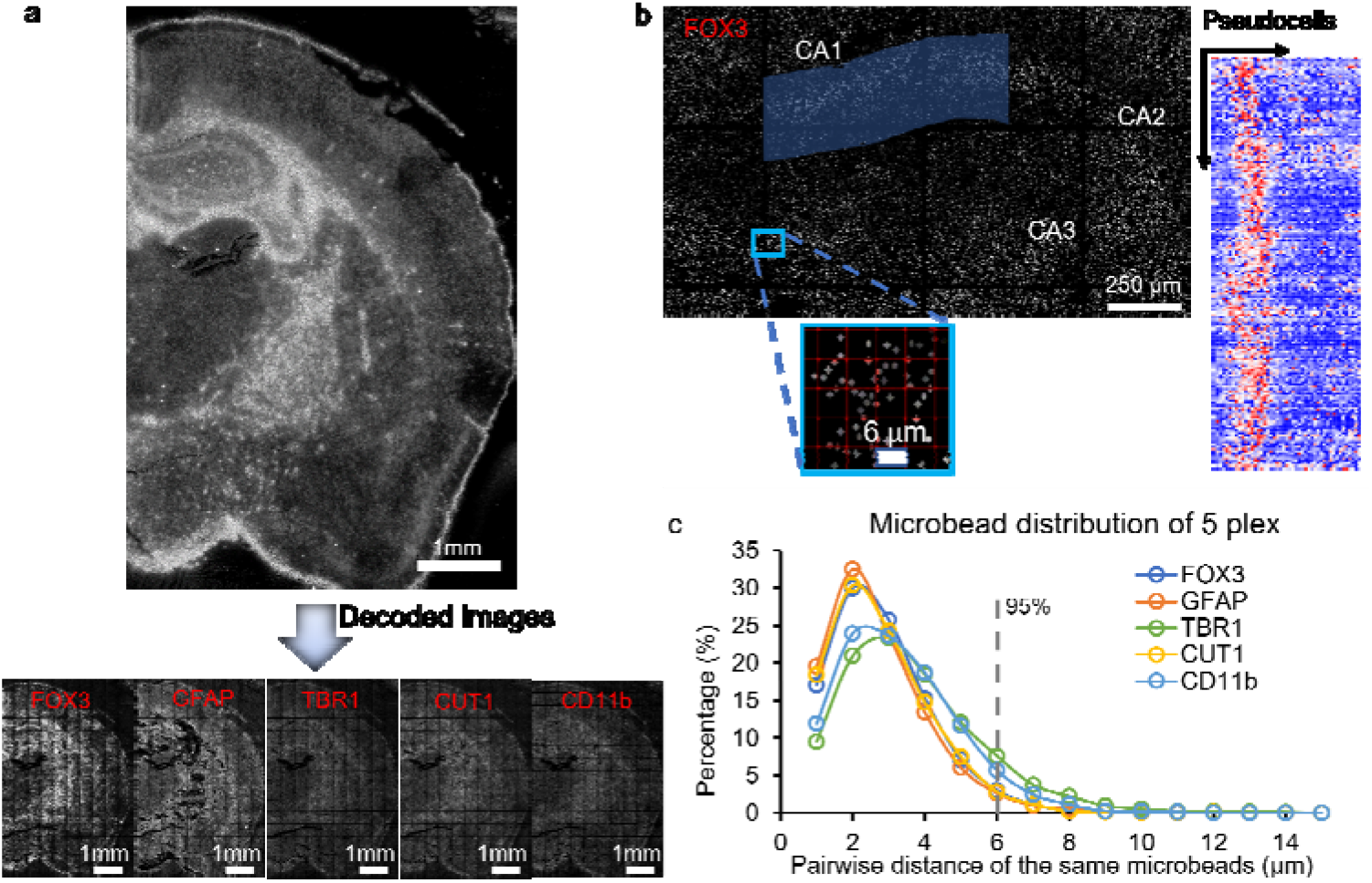
Multiplexed detection of 5 markers by spatial MIST. (a) Image of co-detection of 5 proteins on the MIST array and the decoded images below corresponding to individual proteins. (b) Distribution of FOX3 where only the microbeads for detecting FOX3 are visualized, and all the other areas are masked in dark. The red grid on the zoom-in image defines the boundary of pseudocells and their spa ial addresses. The heatmap on the right corresponds to the fluorescence intensity of the shaded CA1 region on the left image. (c) Distribution of pairwise distance of the same microbeads when 5 types of microbeads are used for multiplexed detection. Dash line indicates the pairwise distance with 95% probability that at least one copy of microbead should be found for any of 5 types.

Measurement of 31 proteins simultaneously for a tissue slice is achievable with the same detection time as 1 or 5-plex detection. At various regions of the brain, the distributions of bright microbeads on the MIST array are very distinctive due to different protein expression (Figure 4a). As expected, the CA1 region has more bright microbeads because of high local density of neuronal cells. We also examined the microbead-microbead distances to determine the size of pseudocells. It was found 15 μm is sufficient to compose a complete protein expression profile with 95% probability, and 20 μm is corresponding to 99% probability (Figure 4b). Most likely due to technical issues, some microbeads have longer pairwise distances. To facilitate downstream analysis, we conservatively selected 20 μm as the pseudocell size to minimize the influence of missing certain proteins in the analysis. These sizes are larger than a typical brain cell at about 10 μm, and they could be optimized in the future by approaches such as detecting two proteins on one microbead instead of one or using smaller size of microbeads. Nevertheless, codetection of 31 proteins quickly within 2 h total time is still ~10 times faster than other similar high-multiplex technologies.

**Figure 4.**
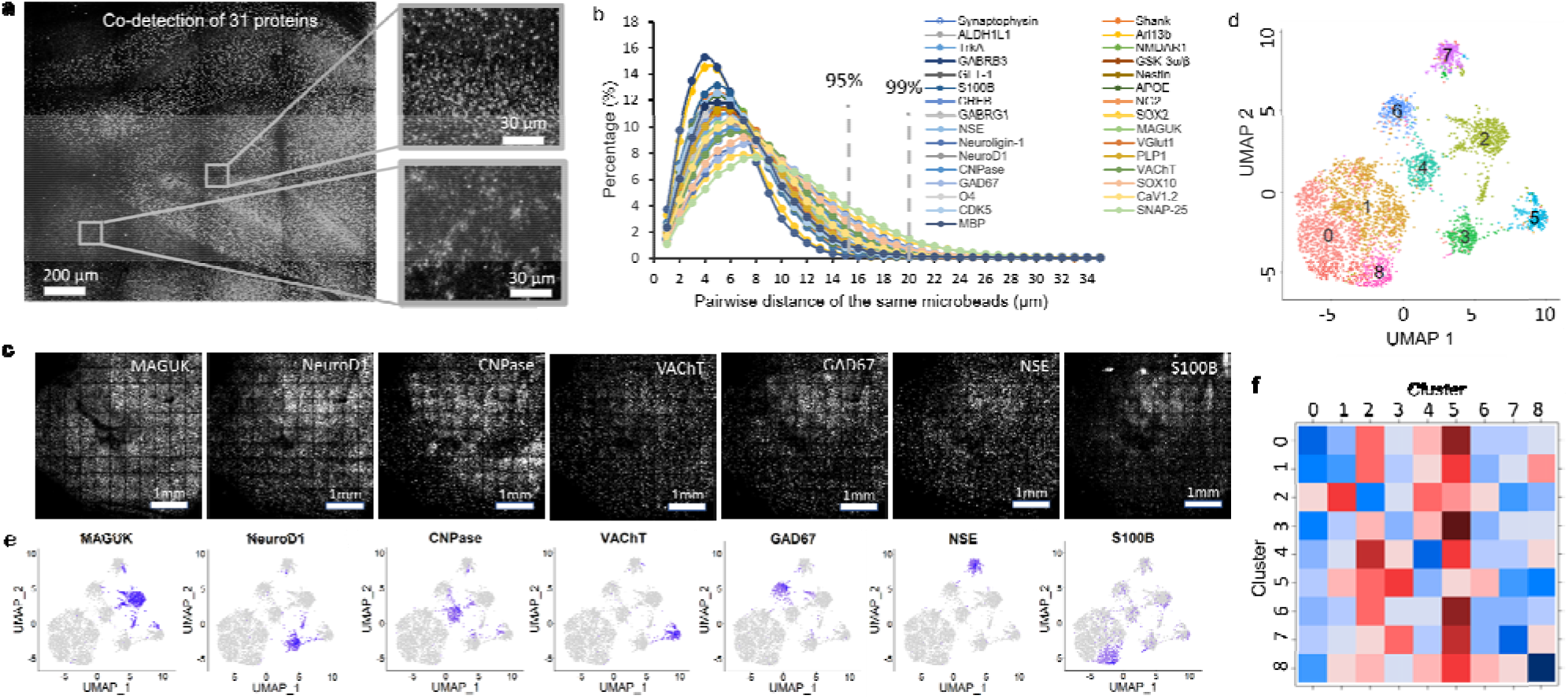
Multiplexed detection of 31 proteins by spatial MIST. (a) Stitched MIST array image of part of mouse brain coronal section with two zoom-in images on the right. (b) Distribution of pairwise distance of the same microbeads when 31 microbeads are used for multiplexed detection. Dash line indicates the pairwise distance with 95% probability that at least one copy of microbead should be found for any of 31 types. (c) Sample MIST array images for individual proteins after decoding. Scale bars represent 1 mm. (d) UMAP clustering of pseudocells where 9 clusters are identified by the algorithm. (e) Location of selected proteins on the UMAP clusters. (f) Heatmap visualization of spatial proximity distance between various clusters generated by UMAP.

The 31-protein panel includes a wide range of surface markers, signaling pathway proteins, and transcription factors that represent various brain cell subtypes, neuron functions and biological processes. The pseudocells were clustered by UMAP in Figure 4d to identify regional variations and their spatial distributions. 9 clusters, or 9 types of pseudocells, are identified that possess distinctive overall protein expression profiles (Figure 4e). Interestingly, 7 proteins including MAGUK, NeuroD, CNPase, VAChT, in any clusters. The corresponding images for these 7 proteins in Figure 4c show they have different distribution across the brain. We further analyzed the similarities between the pseudocell clusters to examine the neighborhood of pseudocells. The similarity was defined as the average minimum nearest neighbor distance between the clusters (Figure 4f). The diagonals on the heatmap represent the average minimum nearest neighbor distance between its own cluster, on the condition that a pseudocell cannot be its own neighbor, which generally has the lowest distances. Cluster 0 has the strongest low-distance values (blue) which has a decent amount of similarity between other clusters, suggesting that the pseudocells within that cluster are more scattered throughout the image. On the contrary, cluster 5, represented by VAChT, is generally far away from any of other clusters, indicating that these pseudocells, or small regions, are relatively independent. The neurotransmitter VAChT is a biomarker for cholinergic neurons that transports vesicular acetylcholine and are distributed in cell bodies, axons and axonal nerve terminals.^33^ It is highly possible that VAChT distinguish itself from the other proteins in our panel by labeling granular vesicles in synapses and axons, so the spatial proximity distance is relatively longer.

## CONCLUSION

In summary, we have developed a spatial MIST technology for rapid, one-shot detection of dozens of proteins markers on a brain slice. The spatial MIST transfers the cleaved barcode DNAs from IF assayed tissue to a high-density microbead array. It exhibits high fidelity and high spatial resolution in detection. Further codetection of 5 markers reveals the distributions of individual markers. We demonstrated the multiplexed detection of 31 proteins simultaneously on a brain slice also within 2 h. Clustering of pseudocells by UMAP found 9 distinctive sub-regions of the brain. Spatial proximity analysis resulted in the discovery of VAChT-rich regions that are distributed away from other sub-regions. With the advantages of rapid multiplexed assay and simplicity, we envision that spatial MIST can find broad biomedical applications in the clinic and mechanistic studies of brain diseases as well as other human diseases as long as tissue section is concerned.

## ASSOCIATED CONTENTS

### Supporting Information

The Supporting Information is available free of charge on the ACS Publications website at DOI: List of antibodies, oligonucleotides and decoding design table.

## AUTHOR INFORMATION

### Corresponding Authors

*

### Notes

The authors declare no competing financial interest.

## ACKNOWLEDGEMENT

This work was supported by the National Institutes of Health R21AG072076 (JW) and R01GM128984 (JW). We also thank Dr. Richard Moffitt for the discussion and data analysis.

## Notes

### Competing Interest Statement

The authors have declared no competing interest.

